# Deep learning image recognition enables efficient genome editing in zebrafish by automated injections

**DOI:** 10.1101/384735

**Authors:** Maria Lorena Cordero-Maldonado, Simon Perathoner, Kees-Jan van der Kolk, Ralf Boland, Ursula Heins-Marroquin, Herman P. Spaink, Annemarie H. Meijer, Alexander D. Crawford, Jan de Sonneville

## Abstract

One of the most popular techniques in zebrafish research is microinjection, as it is a rapid and efficient way to genetically manipulate early developing embryos, and to introduce microbes or tracers at larval stages.

Here we demonstrate the development of a machine learning software that allows for microinjection at a trained target site in zebrafish eggs at unprecedented speed. The software is based on the open-source deep-learning library Inception v3.

In a first step, the software distinguishes wells containing embryos at one-cell stage from wells to be skipped with an accuracy of 93%. A second step was developed to pinpoint the injection site. Deep learning allows to predict this location on average within 42 µm to manually annotated sites. Using a Graphics Processing Unit (GPU), both steps together take less than 100 milliseconds. We first tested our system by injecting a morpholino into the middle of the yolk and found that the automated injection efficiency is as efficient as manual injection (~ 80%). Next, we tested both CRISPR/Cas9 and DNA construct injections into the zygote and obtained a comparable efficiency to that of an experienced experimentalist. Combined with a higher throughput, this results in a higher yield. Hence, the automated injection of CRISPR/Cas9 will allow high-throughput applications to knock out and knock in relevant genes to study their mechanisms or pathways of interest in diverse areas of biomedical research.

## Introduction

Microinjection is one of the most powerful techniques used in zebrafish (*Danio rerio*), as it allows to follow cell fate [1], evaluate pathogenesis of bacteria [2], produce chimeric individuals [3], study tumour progression [4,5], manipulate protein levels [6,7] and create genetically altered lines [8]. The intrinsic biological properties of zebrafish make it particularly amenable to this technique, since these cyprinids are highly fecund, a spawning pair typically producing more than 400 eggs at a time. Moreover, fertilization is external and spawning is confined to a brief period at dawn (natural or artificial), allowing for timing of the experiments. Furthermore, the chorion of zebrafish eggs is supple and easy to pierce.

Classically, injection of tracer dyes is used to identify single cell populations [9,10], to follow cell lineages and to build fate maps in zebrafish [1,11]. The development of molecular methods for the zebrafish model enabled functional studies by manipulating the expression of specific genes. Injection of messenger RNA (mRNA) can be used to overexpress and misexpress a specific protein [12], while morpholino antisense oligonucleotides (MOs) can be employed to knock down a given target gene [13]. Morpholinos are synthetic oligonucleotides designed to be complementary to a specific RNA target. Typically they target the translation start site blocking translation initiation. Alternatively, MOs can also be designed to mask splicing sites, hindering the proper processing of precursor mRNA. In zebrafish mRNA and MO injections are simply performed by introducing a fine-tipped needle into the yolk of one-cell stage eggs and delivering nanoliter volumes of the injection material into it [14]. As cytoplasmic streaming will move the mRNA or MOs into the cytoplasm, it is not necessary that the injection targets the cell. While injection into the yolk requires some skill, it can usually be learned within a few weeks. Nevertheless, injections of mRNAs and MOs have their drawbacks. First of all, the effect is only transient, *i.e.* the injected molecules will be degraded and/or diluted with time. Moreover, in the case of mRNA injection, tissue-specific upregulation is not possible and a given mRNA will be expressed in all tissues indiscriminately. Also, the specificity of MO antisense technology has recently been questioned. Indeed, MOs can sometimes lead to misleading results, in many cases due to toxicity and off-target effects, but also due to the difficulties in estimating their efficacy and controlling their dosage [15]. In a recent study [16], loss-of-function mutations for ten different genes previously thought to have an essential role in development failed to recapitulate the corresponding morpholino-induced phenotypes. In several cases, the discrepancy between mutant and morphant phenotypes, could be explained by genetic compensation mechanisms that occur in mutants [17], however, undoubtedly rigorous controls are required to ascertain the reliability of MO-induced phenotypes [15,18,19].

In the last years, with the implementation of targeted nuclease techniques in the zebrafish, the demand for genetic evidence to define gene function has greatly increased. Fortunately, after a somewhat slow start using zinc-finger nucleases (ZFNs) [20] and transcription activator-like effector nucleases (TALENs) [21], the adaptation of the prokaryotic CRISPR/Cas9 (clustered regularly interspaced short palindromic repeats/CRISPR associated protein 9) defence system to engineer genomes [22] has revolutionized reverse genetics in zebrafish.

The CRISPR/Cas9 method builds on the type II CRISPR system of the adaptive immunity of certain archaea and bacteria [23]. When infected by a bacteriophage, these prokaryotes respond by integrating fragments of the viral genome into clustered regularly interspaced short palindromic repeat (CRISPR) loci. These loci are subsequently transcribed and processed to CRISPR RNAs (crRNAs). crRNAs form a base-paired structure with a transactivating CRISPR RNA (tracrRNA) to recruit CRISPR associated protein 9 (Cas9) [22]. This nuclease is guided to the viral genome by the RNA complex and the target sequence is subsequently cleaved, neutralizing the invading entity. The cleavage site of the Cas9 nuclease is determined by a short sequence adjacent to the target DNA and by sequence complementarity of the crRNA with the target site itself.

Recently the system was adapted and optimised to engineer genomes. crRNA and tracrRNA can be replaced by a single synthetic guide RNA (gRNA) that directs Cas9-mediated cleavage of target DNA [22,24], and the method was implemented in multiple systems including zebrafish [25,26], finally paving the road for knock-ins in this model [27]. Along with *Tol2* mediated transgenesis, a transposon system based on the *Tol2* element of medaka (*Oryzias latipes*) widely used in zebrafish to create transgenic lines [8], the CRISPR/Cas9 system has become an essential tool for genome editing in zebrafish. In this context microinjection is an essential technique. For the creation of genetically altered lines in zebrafish, be it through *Tol2* transgenesis or by means of zinc finger nucleases, TALEN or Cas9 nucleases, it is critical to inject the solution directly into the blastomere at the one-cell stage or at least at the interface between blastomere and yolk [28–31]. Contrary to RNA or MOs, DNA appears not to be transported into the blastomere via cytoplasmic streaming. Moreover, efficiency of all these genome editing techniques is much lower compared to mRNA or MOs injections. Therefore, in order to create genetically altered zebrafish lines it is essential to master microinjections into the cell. This can be challenging as this type of injection requires long training and excellent technical skills.

### Automated microinjection system

The first version of our automated microinjection system featured half-spherical wells, moulded in agarose gel, which allowed for high-throughput microinjection into the yolk of zebrafish eggs. This was used for microinjection of bacteria, morpholinos [32] and cancer cells [33]. The great advantage over other systems was the higher batch size and speed of the injections. As the initial cell division steps in zebrafish embryos occur in intervals of 20-40 minutes, speed is crucial for the accuracy, reproducibility and number of experiments.

In our experience, it is apparent that injections into the middle of the yolk are less suitable for DNA injections. Therefore as a first step, the program “click-to-inject” was developed to test the efficiency of injections closer to the first cell [33]. With this, we noticed that we could achieve a great increase in efficiency, similar to manual injections done into the first cell. Therefore, we set out to automate this procedure.

In this study we demonstrate the results of autonomous site selection and injection for CRISPR/Cas9 and DNA manipulation of the zebrafish genome.

## Materials and methods

### Animals

At the Luxembourg Centre for Systems Biomedicine, wild type adult zebrafish (AB strain) are maintained in the Aquatic Facility according to standard protocols [34]. Zebrafish eggs were obtained by natural spawning on the day of each experiment, kept in 0.3X Danieau’s solution (14 mM NaCl, 2 mM KCl, 0.12 mM MgSO_4_, 1.8 mM Ca(NO_3_)_2_, 1.5 mM HEPES pH 7.5 and 0.03 M methylene blue), and staged by morphology (one-cell stage) for the injections. After each series of injections, the eggs were incubated at 28 °C (±0.5) and evaluated daily until 5 days post-fertilization (dpf). At the Institute of Biology, Leiden University, wild type adult zebrafish (strain AB/TL) are maintained and handled according to standard protocols [34]. Zebrafish eggs were obtained by natural spawning on the day of each experiment, and after each series of injections, the eggs were incubated at 28.5 °C in egg water (60 μg/ml sea salt, Sera Marin, Heinsberg, Germany) and evaluated at 6 hours post-fertilization (hpf), 1 dpf and 5 dpf. Anaesthesia of larvae used for live imaging and COPAS [32,35] analysis was done with 0.02% buffered Tricaine (3-aminobenzoic acid ethyl ester, Sigma-Aldrich) in egg water.

### Ethics statement

All procedures with zebrafish were performed in accordance with European laws, guidelines and policies for animal experimentation, housing and care (European Directive 2010/63/EU on the protection of animals used for scientific purposes). The operation of the Aquatic Facility at the Luxembourg Centre for Systems Biomedicine is allowed by the relevant agencies of the Government of Luxembourg (authorization through the Grand-Ducal decree of 20 January 2016). Maintaining and handling of adult zebrafish at the Institute of Biology, Leiden University, is in compliance with the animal welfare directives of the local animal welfare committee.

### Morpholino antisense oligonucleotide

The translation blocking morpholino for *slc45a2* (solute carrier family 45 member 2) was obtained from Gene Tools according to Dooley *et al*., 2012 [36] with the following sequence: 5’-GCTGGTCCTCAGTAAGAAGAGTCAT-3’. In addition, a 3’ fluorescein modification was included, which allowed fluorescent differentiation of injected eggs. A standard MO with sequence 5’-CCTCTTACCTCAGTTACAATTTATA-3’ was used as an injection control. In both cases, stock solutions (1 mM ~ 8 ng/nL) were prepared according to the specifications of the provider and titrated working solutions were freshly prepared for each experiment.

### Preparation of Cas9 mRNA and *slc45a2* sgRNA

Both *slc45a2* sgRNA and Cas9 mRNA were prepared according to Gagnon *et al*., 2014 [30]. Briefly, the *slc45a2* DNA template was synthetized with T4 DNA polymerase (New England BioLabs) using the oligonucleotides: *slc45a2*-specific (taatacgactcactataGGTTTGGGAACCGGTCTGATgttttagagctagaaatagcaag) and constant (AAAAGCACCGACTCGGTGCCACTTTTTCAAGTTGATAACGGACTAGCCTTATTTTAACTTGCTATTTCTAGCTCTAAAAC). The sgRNA was synthetized using T7 RNA polymerase (Ambion MEGAscript) and then diluted to 400 ng/µl. Cas9 mRNA was synthetized using the pCS2-Cas9 plasmid [37], transcribed using the SP6 mMessage mMachine kit (Ambion) and finally diluted to 600 ng/µl.

### Manual microinjections of *slc45a2*-MO and *slc45a2* sgRNA/Cas9

Manual microinjections of zebrafish embryos were done as described by Rosen *et al*. 2009 [14]. Briefly, gene knockdown was achieved through manual microinjection of 2 nL of *slc45a2*-MO into the center of the yolk of wild-type zebrafish embryos at one-to two-cell stage using an Eppendorf FemtoJet 4X® microinjector. We titrated the amount of the *slc45a2*-MO to 3.7 ng per injection, and the same amount was used for the control MO. On the other hand, gene editing was achieved through manual microinjection of 4 nL of *slc45a2*-sgRNA, Cas9 and phenol red in a single mix into the cell cytoplasm of wild-type zebrafish embryos at one-cell stage using an Eppendorf FemtoJet 4X® microinjector. After each series of microinjections, the eggs were incubated at 28 °C (±0.5) in 0.3X Danieau’s medium and evaluated daily until 5 dpf to record non-viable embryos, (*i.e.* non-fertilized, fluorescent negative at 6 hpf for MO-injections, dead and dysmorphic embryos/larvae from 1 to 5 dpf), and the efficiency of injection displayed by the albino phenotype.

### Manual microinjection of DNA

Manual microinjection of zebrafish embryos was done as described previously using standard methods [38] using an Eppendorf FemtoJet microinjector. Briefly, a DNA construct containing a GFP fusion gene expressed under a constitutive promoter (*actb*:-NLSmCherry-IRES-GFP) in the Gateway *Tol2* vector was co-injected together with *Tol2* transposase RNA in a volume of 1 nL at a concentration of 25 ng/μL for both the DNA construct and the *Tol2* transposase RNA. Injections were performed at the one-cell stage at the blastomere/yolk boundary. After each series of injections, the eggs were incubated at 28.5 °C in egg water and evaluated at 6 hpf and 1 dpf to record non-viable embryos, and at 5 dpf for fluorescent signal.

### Automated microinjection of *slc45a2*-MO

Automated microinjection of zebrafish embryos using the robotic injector (Life Science Methods BV) was done following guidelines described in Spaink *et al*. 2013 [33]. Briefly, gene knockdown was achieved through fully automated microinjection of 2 nL of *slc45a2*-MO (3.7 ng per injection) into the yolk of wild-type zebrafish embryos at one-to two-cell stage. A 1% agarose covered grid (9 blocks x 100 wells) was used to carefully arrange the zebrafish embryos, and then placed in the motorized stage coupled to a controlled and motorized micro-manipulator. The needle that was first loaded with the sample and calibrated for a microinjection volume of 2 nL was then placed into the micro-manipulator (Eppendorf FemtoJet). After the robotic injector was properly set (position of grid and needle) automated injection occurred at a speed of approximately 100 embryos per 160 seconds (1 block in the grid). All components of the robotic injector are connected to a controlling computer that is equipped with a software control program written in Python. After each series of microinjections, the eggs were incubated at 28 °C (±0.5) in 0.3X Danieau’s medium and evaluated daily until 5 dpf to record non-viable embryos, (*i.e.* non-fertilized and fluorescent negative at 6 hpf, dead and dysmorphic embryos/larvae from 1 to 5 dpf), and the efficiency of injection displayed by the albino phenotype.

### Automated microinjection of *slc45a2* sgRNA/Cas9

Automated microinjection of zebrafish embryos using the robotic injector (Life Science Methods BV) was done following guidelines described in Spaink *et al*. 2013 [33]. Briefly, microinjection of 4 nL of *slc45a2*-sgRNA, Cas9 and phenol red in a single mix was done in zebrafish embryos at one-cell stage. A 1% agarose covered grid (9 blocks x 100 wells) was used to carefully arrange and orient one by one the zebrafish embryos so that the first cell is visible. The grid was then placed in the motorized stage coupled to a controlled and motorized micro-manipulator. The needle that was first loaded with the sample and calibrated for a microinjection volume of 4 nL was then placed into the micro-manipulator (Eppendorf FemtoJet). After the robotic injector was properly set (position of grid and needle) the automated cell injections took place at a speed of approximately 100 embryos per 2 minutes and 30 seconds (1 block in the grid). All components of the robotic injector are connected to a controlling computer that is equipped with a software control program written in Python. This allowed to have the total count of the type of microinjections performed depending on the image classification (see section below), *i.e. “*Close-to-cell” #72, “Center” #13, “Empty” #1, “Two-Cell” #14, and “Sick” #0, resulting in 100 injected eggs. After each series of microinjections, the eggs were incubated at 28 °C (±0.5) in 0.3X Danieau’s medium and evaluated daily until 5 dpf to record non-viable embryos, (*i.e.* non-fertilized at 6 hpf, dead and dysmorphic embryos/larvae from 1 to 5 dpf), and the efficiency of injection displayed by the albino phenotype.

### Deep learning algorithm for image classification

As a first step we used the Inception v3 network to learn to distinguish between five different categories: “Empty”, “No-Cell”, “First-Cell”, “Two-Cell”, “Sick” (this term is used to refer to non-viable eggs). We used a total of 11,000 annotated images. To prevent overfitting, we artificially increased the number of training samples by performing four types of image transformation: *1)* rotations about the center of the image; *2)* zooming by a factor 0.9-1.1; *3)* shifting by 28 pixels orthogonally in the +/- x and y direction, and *4)* flipping the image horizontally. The neural network architecture consisted of: *1)* the top part of the Inception v3 network (containing all inception blocks); *2)* a 2D global spatial average pooling layer; *3)* a fully connected layer of 1024 nodes with ReLU activation function, and *4)* a fully connected layer of 5 nodes, with softmax activation function. Training of the classification step was done using the Adam stochastic optimizer [39], with a learning rate of 10^-4^. For a more in-depth description see the supporting information (S1).

### Deep learning algorithm for finding the injection site

For the injection point determination, we translated the (x y) coordinates to a vector in a triangular mesh using a barycentric coordinate system. We let the outputs of the neural net correspond to vertices in the mesh. In our case, we used 160 vertices.

The neural network architecture consists of: *1)* the top part of the Inception v3 network (containing all inception blocks); *2)* a 2D global spatial average pooling layer; *3)* a fully connected layer of 1024 nodes with ReLU activation function, and *4)* a fully connected layer of 160 nodes, with softmax activation function. We used 2724 images for training and 674 images for validation (these are the same images as used for label “first-cell” in the classification step). Training of the injection point determination step was done using the Adam optimizer, with a learning rate varying from 10^-3^ to 10^-5^. More details can be found in the supporting information (S1).

### Software and hardware

For deep learning and robot control we used a Shuttle SZ170R8 equipped with an Intel Core i3 6100 CPU, 16 GB kit Kingston DDR4 2133Mhz, ECC memory and an NVidia GeForce GTX 1070 GPU. Installed software are: Keras 1.2.2, Theano 0.9.0, NumPy 1.11.0, SciPy 0.17.0. For the analysis of the data, raw data for all the series of microinjections was processed in excel. Statistical analysis was done using excel and GraphPad Prism 6 followed by unpaired *t*-test with Welch’s correction for single comparisons (when applicable). The criterion for statistical significance was P<0.05. Graphs were plotted using GraphPad Prism 6 and error bars on all graphs represent standard deviation.

### Microscopy and fluorescent analysis

At the Institute of Biology, Leiden University, representative pictures were taken using a Leica M205 FA stereo fluorescence microscope equipped with a DFC345 FX monochrome camera. Fluorescent signal was quantified using a Complex Object Parameter Analyzer and Sorter (COPAS, Union Biometrica). At the Luxembourg Centre for Systems Biomedicine, fluorescent sorting of fluorescein positive embryos (for *slc45a2*-MO injections) was done using a Nikon SMZ25 stereomicroscope. Representative pictures of control larvae and injected larvae displaying an albino phenotype were taken using the Nikon SMZ25 stereomicroscope equipped with a Nikon Digital Sight DS-Ri1 camera.

## Results and discussion

### Manual and automated injections of *slc45a2*-MO

In order to test MO efficiency of manual and automated injections we employed a translation-blocking MO against *slc45a2* (solute carrier family 45 member 2). Downregulation of this gene induces albino and/or hypo-pigmented morphants, as the melanophores are unable to produce melanin [36]. Manual and automated yolk microinjections were performed in parallel, and in both cases the induced albino phenotype was assessed in larvae at 3 dpf. The results obtained with both microinjection approaches are comparable and show that downregulation of *slc45a2* is highly efficient using morpholino antisense technology (Fig 1A). Additionally, the manual injections were performed by two different experimentalists (Fig 1B) and this shows that efficiency and variation of efficiency obtained by manual morpholino injections differs from person to person and, surprisingly, the variation of the efficiency of the automated injections is slightly larger.

**Fig 1.**
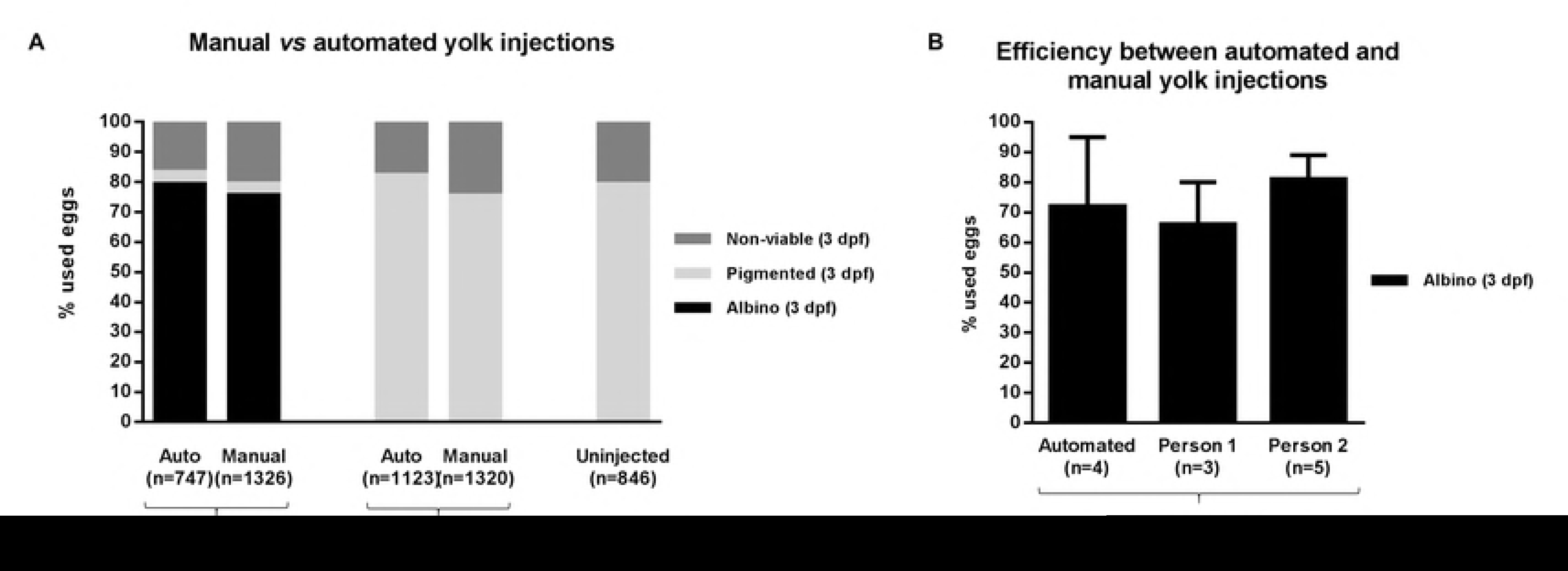
Morpholino knockdown efficiency with manual and automated injections. **A.** The survival and knockdown efficiency of *slc45a2*-MO manual and automated (auto) microinjections were measured as the number of larvae displaying an albino phenotype at 3 days post-fertilization (dpf). Control-MO injected larvae and uninjected larvae were processed in parallel and the resulting pigmented (wild-type) larvae were also counted at 3 dpf. “n=” indicates the number of eggs used to obtain this cumulative result. **B.** Efficiency comparison between the automated injection into the yolk and manual injections performed by two independent experimentalists (not statistically significant). “n=” indicates the number of experiments used to calculate the average and standard deviation.

### Semi-automated “click-to-inject”

After demonstrating that automated injection into the yolk is an efficient way to generate morphants, we sought to apply the robotic injector for generating CRISPR/Cas9 mutants for *slc45a2*. To investigate the dependence on the injection location we used the “click-to-inject” program [33] to test the efficiency of injections closer to the first cell. In the “click-to-inject” program the injection depth is set, but the (x y) position is chosen by the operator. To inject, the operator moves the mouse pointer to a specific site (*e.g.* the first cell) and clicks to trigger an injection and a subsequent movement to the next egg. Based on this, next we set out to develop an automated image recognition to more precisely identify the first cell and to automate CRISPR/Cas9 injections.

### Imaging conditions

In manual microinjection setups, as well as in standard microscopy, near-perfect imaging conditions are applied with lighting from the bottom and imaging from the top, or vice versa. As the zebrafish egg is very transparent, epi illumination from below is not suitable; most contrast and edges are then lost. As the egg is spherical, a ring-light displays a very bright circle on top of the egg. Therefore, to obtain better and more reproducible imaging conditions in different locations, we placed a large (L x B = 60 x 80 cm^2^) diffuse light source above the robotic injector. Five different classes were used to annotate the images (Fig 2). In the “Inject” class the ideal injection position for automated microinjection is also annotated. Instead of injecting directly into the zygote, we have chosen to inject in the yolk, close to the visible zygote. The reason is that injections in a thin zygote (less ideal orientation, or very early stage) would often cause a rotation of the egg, and bounce the needle off. Injections into the yolk-blastomere boundary almost never show this problem, and thus gave a higher yield.

**Fig 2.**
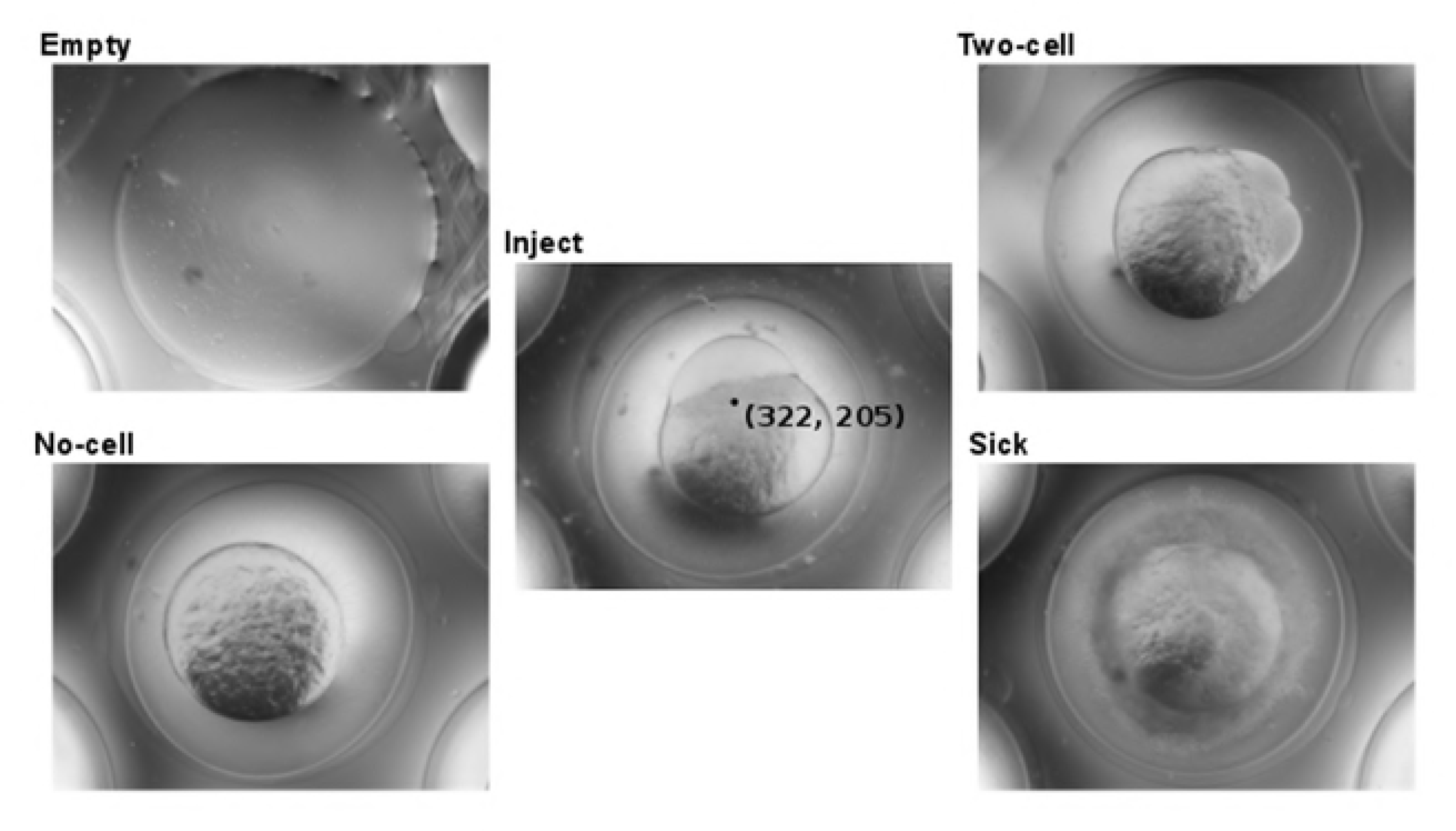
Imaging classification for injection. Representative digital images measured from below of an agarose grid (“Empty”) that supports zebrafish eggs with the first cell visible (“Inject”) or not visible (“No cell”), eggs in a two- or higher cell stage (“Two cell”) or non-viable eggs (“Sick”). In the “Inject” image an injection location is indicated by a black dot with (x y) coordinates.

### Machine learning

Initially, we tried a classic approach of machine vision on these images. The Hough circle transform [40] allowed us to detect the yolk with an above 90% accuracy (data not shown). However, the next step to find the first cell was problematic. In cases where the shadow of the micromanipulator overlapped with the first cell, the edge detection algorithm failed. As an alternative to edge detection, we annotated a database of images with injection positions. We used a Fast Fourier Transform (FFT) algorithm to find a closest matching egg in this database and used that image to infer an injection position. This worked reasonably well with a peak error (distance between calculated position and annotated position) of 20 µm. However, when a good match could not be found, the error was quite high, and as a result the tail of the error was quite large (data not shown). An explanation for the large variation in results is that there is also a large variation in first cell shapes, especially when looking from an arbitrary angle. It can be an early very thin line up to about a third of the yolk depending on the developmental stage and orientation of the egg. To overcome this variation, we could make the annotated database larger, to increase the chance of a close match. Nevertheless, the downside of this solution is that more images have to be compared, and this takes more processing time during injection. Thus, we sought to apply a better approach based on deep learning.

Using a database of annotated images as input, one can also train a deep learning network. Instead of comparing images during runtime, one trains an algorithm that is afterwards used to interpret new images. The execution time of this algorithm is independent of the size of the training image set. Thus, roughly speaking, the larger the number of annotated images, the higher the accuracy of the algorithm. We used the Inception v3 open source deep learning software [41]. This software has been built and tested to categorize images, based on a large training database of images, initially for the annual ImageNet Large Scale Visual Recognition Challenge (ILSVRC; www.image-net.org). The Inception v3 architecture uses a neural network that takes the pixels of images as input and extracts features. Many features are subsequently built on top of features, in different layers of neurons, in higher and higher levels of abstraction, until the neurons reach an output of defined categories [42]. One advantage of the Inception v3 software is that one can reuse the first layers of feature extraction for a different set of images. This is built on the idea that the basic features, *e.g.* lines and simple patterns, can be used in all higher-level features that are used to train new categories with new sets of images. Within eight hours of training time we reached a 93% accuracy, with an execution time in the order of tens of milliseconds.

After finding the images with a visible first cell, the next step was to determine the injection location. To enable the use of deep learning for this problem, we had to modify the output from categories into an ideal location. When just the pixel (x y) coordinate is used as output, only one pixel of the whole image is correct. With this output the neuronal network cannot easily distinguish between locations closer to the annotated location and further away, and this makes learning impossible. Therefore, we translated the (x y) coordinates to a barycentric coordinate system [43]. The Greek word “*barys”* means heavy and refers to the centre of gravity. In a barycentric coordinate system a grid of triangles is used, with a weight assigned to each vertex. This is used as follows. A chosen grid of triangles is placed on top of each image. The annotated injection position will fall within one triangle; then the weights of these triangle vertices are given a value according to the location within that triangle. These weights sum up to one, whereas the other vertices in the grid are all zero. This vertices output vector then represents the ideal outcome. The advantage is now that a small deviation from this ideal output vector can be scored gradually instead of binary. This then allows for efficient training. A more detailed explanation is available in the supporting information (S1). After eight hours of training we created a table of (x y) coordinates using validation images. We calculated the distance between the annotated injection position and the position as predicted by the deep learning network (Fig 3). The average distance is 42 µm, as depicted in Fig 3B, and for 83% of the images this distance is smaller than 60 µm.

**Fig 3.**
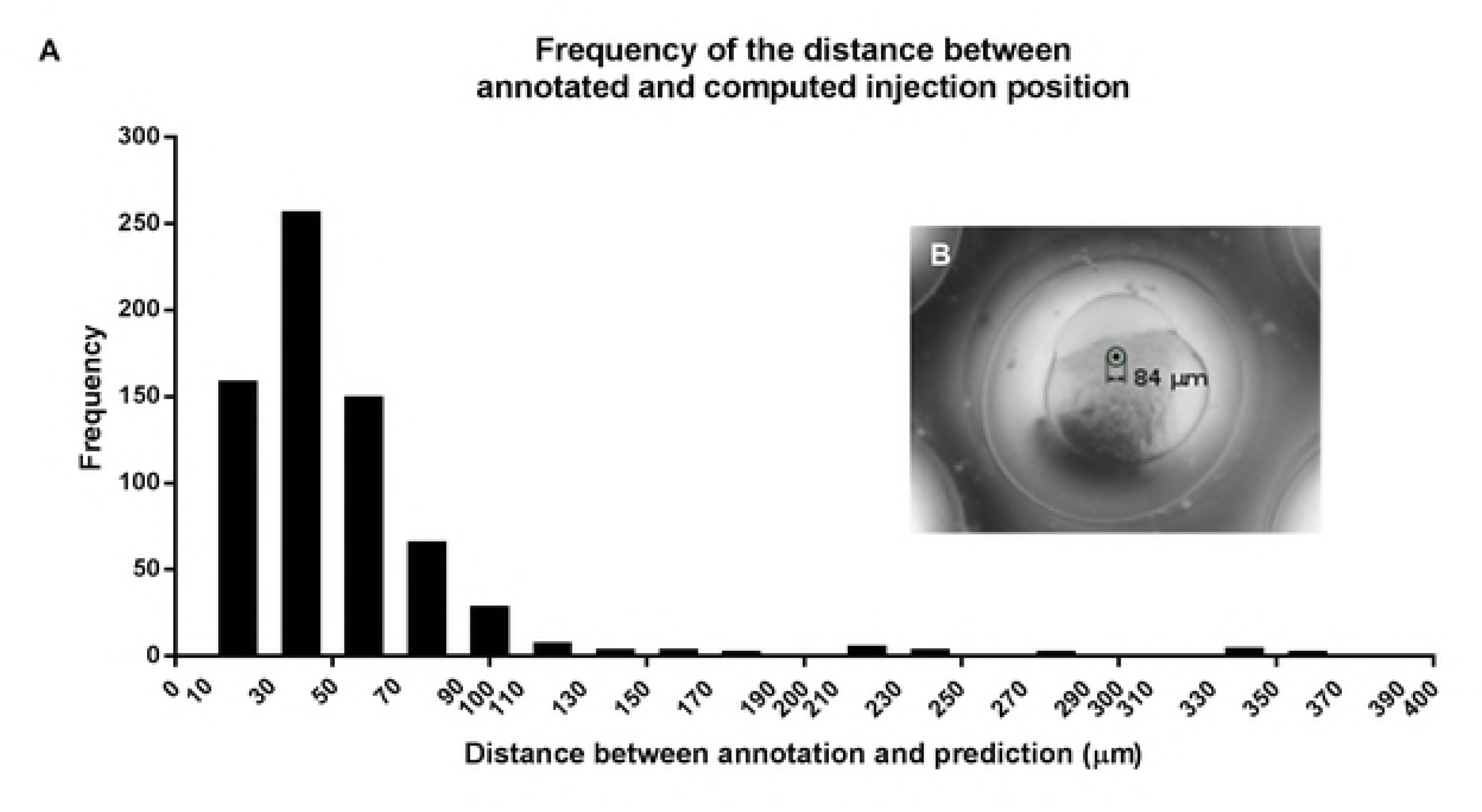
Distance between annotated and computed injection location. **A.** Bar graph depicting the frequency of the distance between annotated and computed injection position (prediction). **B.** Digital image with a circle around an annotated injection point to illustrate the average distance between annotation and prediction.

### Automated injection of *slc45a2* gRNA/Cas9

Trial and error in many laboratories have led to a best practice of injecting into the first cell for the application of the CRISPR/Cas9 editing technique. In our robotic microinjection system, injecting in the middle of the yolk gives the highest speed. Image recognition used to customise an injection location takes time but can increase the injection efficiency. To balance efficiency and speed, and to be able to monitor improvements of our image recognition model, we started by measuring efficiency of CRISPR/Cas9 injections performed in the yolk. Both manual and automated yolk injections gave a very low efficiency of 12% (Fig 4A). Then, with the “click-to-inject” program, resulting in injections closer to the first cell we could generate albino larvae at an almost three times higher efficiency than with the injections in the middle of the yolk (Fig 4A, Semi-auto). Next, using deep learning, we could automate this procedure and with this we reached a slightly lower efficiency when comparing it the “click-to-inject” injections but a higher efficiency than the one obtained with automated and manual injections in the middle of the yolk (Fig 4A, Auto). Still, manual injections into the first cell reached the highest efficiency of 43% (Fig 4A, Manual). Fig 4B shows that both the efficiency and the variation between experiments differs considerably depending on the experimentalist (displayed as P1, P2 and P3). In contrast, here, the automated injections show relatively little variation, also when compare them to the “click-to-inject” injections (Fig 4B, Semi-auto). Also, it can be seen that the efficiency is quickly surpassed by humans given enough experience (P1 and P2). This lower efficiency achieved with the robot can be explained by the injection location – close-to-cell instead of into the zygote – and by the fact that not all the eggs are oriented with a cell visible on the side, despite the fact that they are oriented in the agar grid. Hence, the automated injections tend to be a mixture of injections into the middle of the yolk, and close to the first cell, when the first cell was detected. With this we obtained an efficiency of 24% on average (Fig 4B).

**Fig 4.**
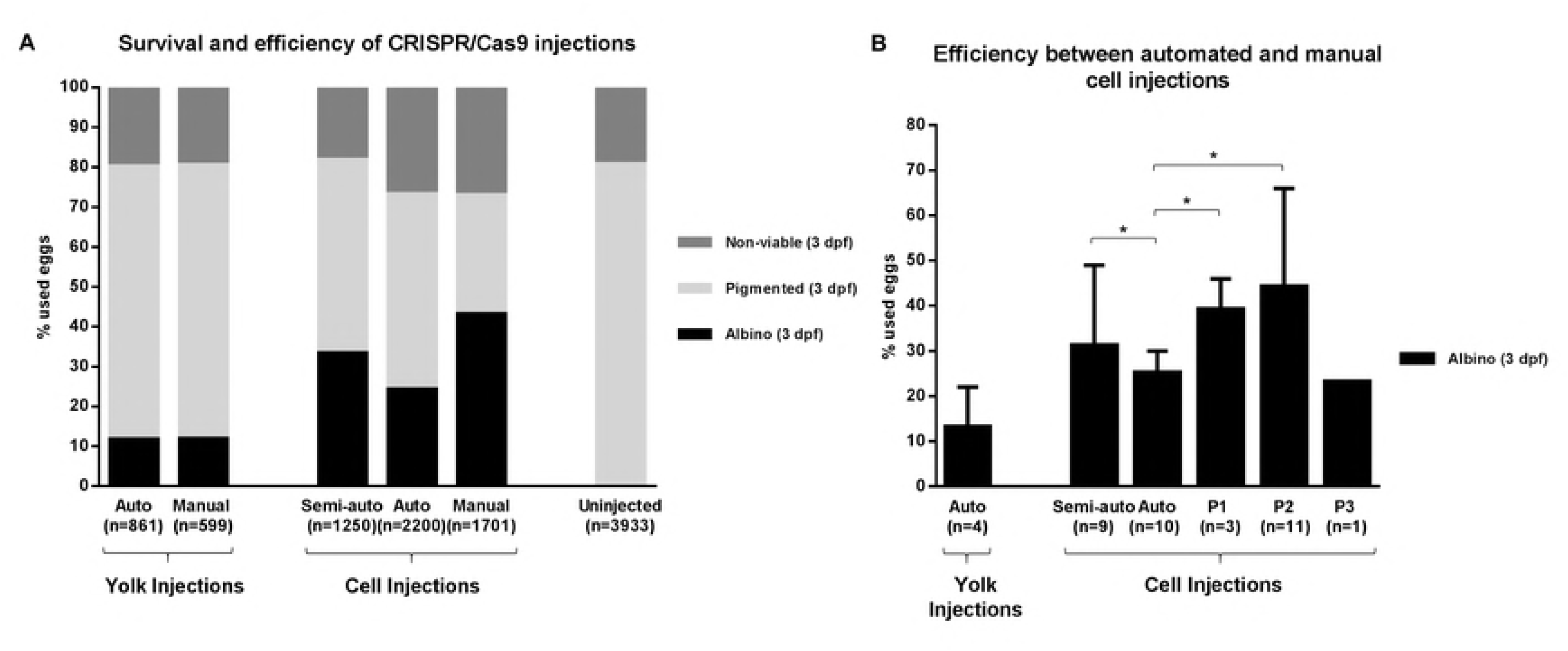
Automated injections of CRISPR/Cas9. **A.** Survival and average efficiency of *slc45a2* gRNA/Cas9 manual, click-to-inject (semi-auto) and automated (auto) microinjections both in the yolk and in the cell were measured as the number of larvae displaying an albino phenotype at 3 days post-fertilization (dpf). Uninjected larvae were processed as controls and the resulting pigmented (wild-type) larvae were also counted at 3 dpf. “n=” indicates the number of eggs that were used to obtain the cumulative results. **B.** Comparison of the average efficiency and standard deviation between the automated (auto), click-to-inject (semi-auto) and manual injections performed by three independent experimentalists (P1, P2, P3). “n=” indicates the number of experiments that were used to calculate the average and standard deviation. * P<0.05.

### Automated injection of DNA

The injections with DNA were performed with an identical robotic microinjection system, but at a different laboratory. Here, we used a COPAS (Complex Object Parameter Analyzer and Sorter) system to measure the efficiency of the injections (Fig 5A). For this, we first measured the highest red fluorescence signal of the uninjected control larvae and took the highest signal as a threshold at 5 dpf. Then we measured the DNA-injected larvae and counted the larvae that passed this threshold. The survival was measured at 1 dpf to focus on differences as a result of the injection. Prior to placing the larvae into the COPAS system, larvae with visible developmental defects were removed. Both the manual and automated injected eggs had a similar relative number of malformed embryos (4% on average, results not shown).

**Fig 5.**
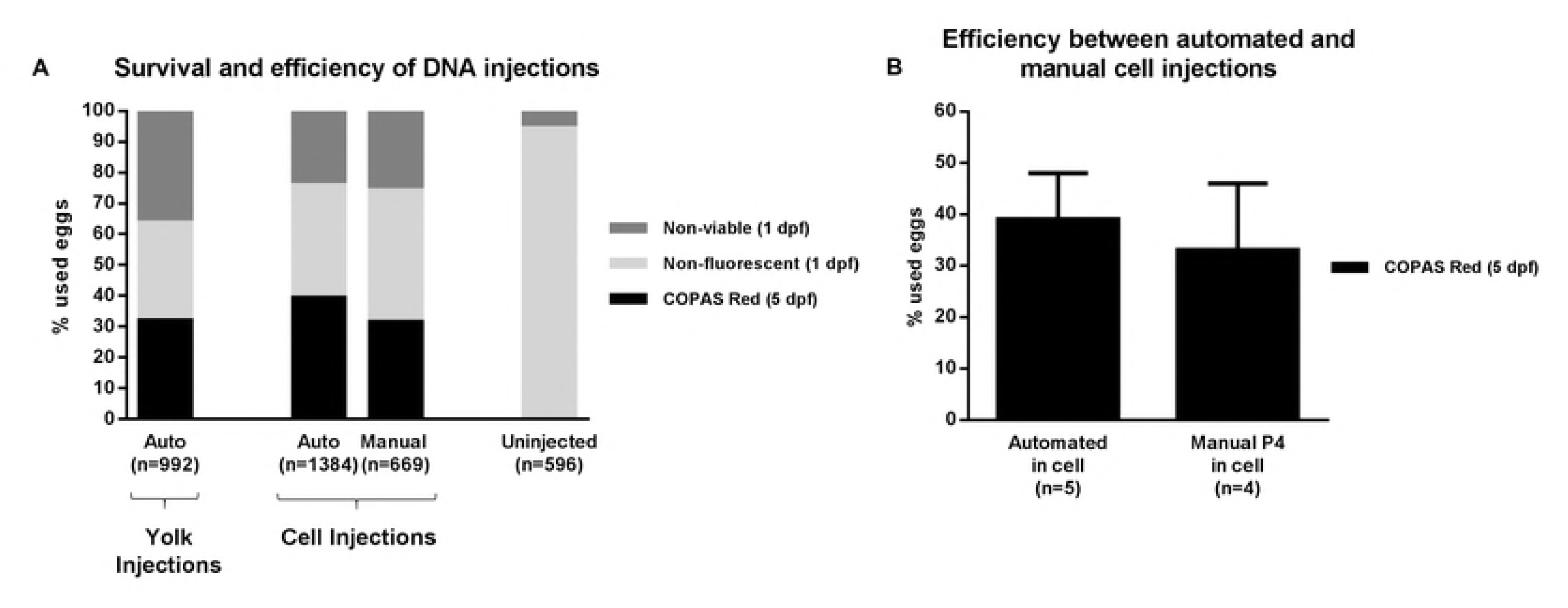
Automated injections of DNA. Average survival and efficiency of DNA automated (auto) and manual injections as measured by the COPAS system. “n=” indicates the number of eggs that were used to obtain the cumulative results. **B.** Comparison of the average efficiency and standard deviation between the automated and manual cell injections. P4 indicates a different experimentalist and “n=” indicates the number of experiments that were used to calculate the average and standard deviation.

These results show that DNA injections are less demanding in terms of injection location. Injections into the middle of the yolk reached an average efficiency of 32%. This can be improved by injecting close to the first cell, when possible, to reach an efficiency of 39%. Surprisingly, here manual injections close to the first cell (personal preference) had a lower efficiency than could be obtained by automated injections and gave on average the same efficiency as injections into the middle of the yolk.

### Microinjection throughput

To calculate the microinjection throughput, we divided the average injection time by the average efficiency. This results in the average time needed for one successfully injected larva. We measured and compared the throughput for the different genetic modifications and experimental setups described in this article, *i.e.* automated and manual injections for gene knockdown by morpholino antisense, gene knockout by CRISPR/Cas9 and transgenesis by *Tol2* (Fig 6).

**Fig 6.**
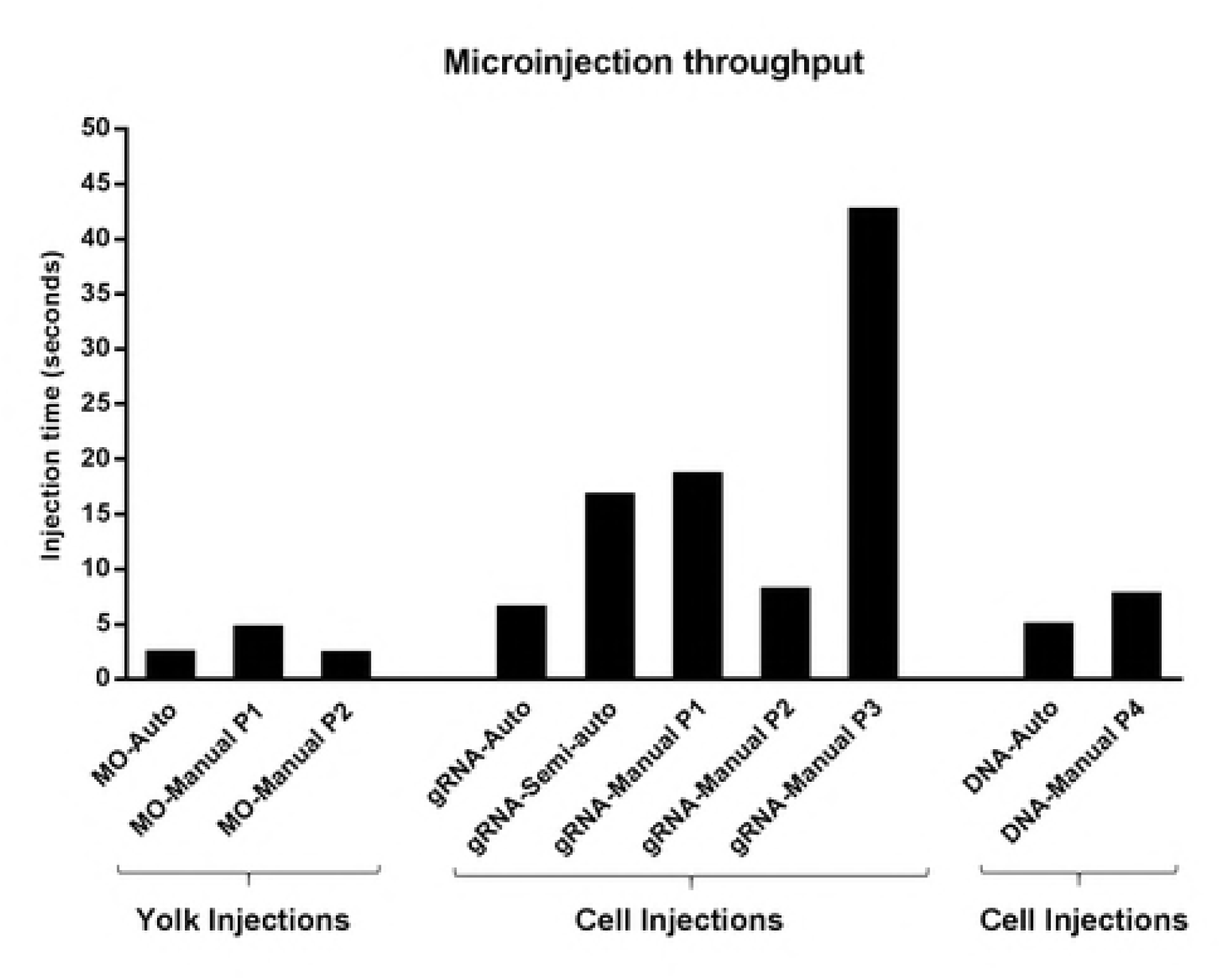
Average injection time required to obtain one positive genetically modified larva. Abbreviations: MO, *slc45a2* morpholino; gRNA, *slc45a2* gRNA/Cas9; DNA, *Tol2* construct; Auto, automated injections; P1-4, four different experimentalists.

In the case of the manual injections, the throughput differs greatly depending on the experimentalist, as experience can lead to a higher throughput by increasing both the efficiency and speed of the injection process. It can also be seen that the robot is on par with fast human performance in case of the morpholino injections, but 1.5 times as fast as average human performance.

With deep learning, a robot can outperform humans on the more complex cell injections. With CRISPR/Cas9 the robot needs 6 seconds of injection time to obtain a positive larva, and humans need 8 up to 43 seconds. On average, the robot is more than three times (3.6x) faster. Manual injections of DNA constructs close to the cell are faster to perform than injections into the cell (2.5 seconds *vs* 6.8 seconds). However, this also reduces the manual efficiency, resulting in a 1.5 times higher throughput of the robot. A movie showing the robotic injection process in real-time is available in the supporting information (S2). The movie shows that the time between capturing the image and placing the cross (demonstrating the calculated injection location) is only about 100 milliseconds.

### Efficiency dependence on the injection location

Contrary to what might be expected, the efficiency of injections into the middle of the yolk to alter the genome were not negligible as the efficiency was 12% for CRISPR/Cas9, 32% for DNA injections and 80% for morpholino injections. Using the measured efficiencies and statistics of image classification we can calculate the efficiencies of injections close to the first cell. During the injections of CRISPR/Cas9, on average 65% of the eggs were oriented with a first cell visible, and 35% were injected into the middle of the yolk. The increase in efficiency, 24%, was caused by 65% of the eggs being injected with efficiency much higher than 12%. Using the efficiency of the yolk injections we can predict the efficiency of injections close to the first cell. Solving the equation 0.65*X+0.35*0.12 = 0.24 for X results in an efficiency of around 30% for injections close to the first cell. For DNA injections we have chosen to not orient the eggs after placing them in a grid, and therefore less eggs, 46%, were injected close to the first cell. Solving the equation 0.46*X + 0.54*0.32 = 0.39 for X results in a predicted efficiency of 47% for injections close to the first cell. Surprisingly, this is much higher than what was obtained by manual injections close to the first cell. These measured and calculated efficiencies can also be used to make a prediction of positive embryos, directly after the injection.

## Conclusion and perspectives

In this study we have demonstrated how we improved an automated injection robot to inject close to the first cell using image recognition in order to enable efficient genome editing in zebrafish embryos. This was accomplished using a modified open-source deep-learning software and annotation of thousands of images. A step-by-step approach of first testing an annotation strategy and efficiency helped to predict the increase in efficiency that can be obtained. Initially we tested the efficiency with a semi-automated click-to-inject program. This click-to-inject approach is also suitable as a first step for other microinjection applications, such as injections into older zebrafish larvae or different organisms.

Because of its transparency, rapid development and easy genetic manipulation, zebrafish have become a key vertebrate model organism for the elucidation of developmental processes. With the advent of CRISPR/Cas9 technology, zebrafish are becoming an even more powerful tool for the study of diverse human disorders. The CRISPR/Cas9 system achieves mutagenesis rate of around 80% for generation of knockout lines [25], and has proven to have fewer side effects than other genome editing technologies. However, generation of specific hereditable mutations or epitope tagging of chromosomal genes in zebrafish is still challenging. Unfortunately, genome editing in zebrafish is unpredictable and efficiency sometimes drops to 3.5% [44]. Therefore, higher number of eggs should be injected for the generation of the expected mutation. Creation of zebrafish mutant lines using CRISPR/Cas9 requires precise injections into or close to the zygote. These types of injections take time to master and are tedious if many batches of hundreds of eggs have to be injected, particularly for the generation of knock-in lines. Our results have showed that efficiency and reproducibility of manual cell injections highly depend on the training stage of the person performing the experiment, making it more difficult to have this technique as a routine procedure in the laboratory. Here, we show the establishment of automated injections as a reliable tool for the generation of CRISPR/Cas9 mutants. This method could also be used for high-throughput gene overexpression studies by microinjection of mRNA. Automated microinjections are simple to learn and allow the cell injection of 100 embryos in 2.5 minutes with comparable efficiency to manual cell injections.

### The need for high-throughput genome manipulation

To date there have been almost 9,000 morpholinos used in zebrafish research. In addition, the adaption of CRISPR/Cas9 editing technology is progressing faster than any other gene silencing method, and even faster than the adoption of morpholino knockdown technology (statistics on zfin.org). However, injections of mRNAs or DNA must be more precise and are more time consuming. Therefore injection can be a limiting step for high-throughput genetic studies. For the moment, there are about 30,000 known gene loci that could be interesting to manipulate in order to investigate their function in development, disease or expressed phenotype (zfin.org). Multiplied with 300 injections that are typically used to obtain a mutant, and multiple mutants per gene, this brings us to tens of millions of injections. Much time and efforts would be saved if this tedious but needed task can be performed mostly by robotic systems.

## Acknowledgements

We would like to thank Dr. Uwe Irion for sharing materials and for his valuable input. We are also grateful to the fish caretaker teams from both Leiden University and the Luxembourg Centre for Systems Biomedicine for their daily valuable work. RB, AHM, and HPS were supported by the Netherlands Organisation for Scientific Research (NWO) Domain Applied and Engineering Sciences (TTW project 13259).

## Supporting information

S1: Deep learning supplement

S2: Movie demonstrating robotic injections with deep learning

S3: Representative images of slc45a2-MO and slc45a2 gRNA/Cas9 S4: Representative images of DNA injected larvae

